# Reciprocalspaceship: A Python Library for Crystallographic Data Analysis

**DOI:** 10.1101/2021.02.03.429617

**Authors:** Jack B. Greisman, Kevin M. Dalton, Doeke R. Hekstra

## Abstract

X-ray crystallography is an invaluable technique for studying the atomic structure of macromolecules. Much of crystallography’s success is due to the software packages developed to enable the automated processing of diffraction data. However, the analysis of unconventional diffraction experiments can still pose significant challenges—many existing programs are closed-source, sparsely documented, or are challenging to integrate with modern libraries for scientific computing and machine learning. Here we describe `reciprocalspaceship`, a Python library for exploring reciprocal space. It provides a tabular representation for reflection data from diffraction experiments that extends the widely-used pandas library with built-in methods for handling space group, unit cell, and symmetry-based operations. As we illustrate, this library facilitates new modes of exploratory data analysis while supporting the prototyping, development, and release of new methods.

## 1 Introduction

The analysis of most diffraction experiments begins with processing diffraction images and ends with refining an atomic model that is consistent with the observed data. Numerous software suites and commandline applications address different stages of the processing pipeline, and these diverse programs are typically combined in order to address the challenges of a particular data set [1, 2, 3, 4, 5, 6, 7]. However, many unconventional diffraction experiments do not fit easily into the processing pipelines established within existing crystallography software. Such experiments often require custom scripts and programs to analyze the resulting data. Recent examples of such experiments include time-resolved pump-probe experiments that investigate the structural dynamics within room-temperature crystals [8, 9]. New software is needed to support custom analyses to improve the development, reproducibility, and adoption of less routine diffraction experiments.

A software library to support such experiments must provide built-in methods to handle space group, unit cell, and symmetry-based operations. This requirement is already met by several general-purpose libraries, such as the `Computational Crystallography Toolbox` (`CCTBX`) and `GEMMI` [4, 10]. However, it is also desirable to facilitate the exploratory inspection of reflection data and to support seamless integration with existing scientific computing software. These additional requirements lower the barrier to implement and test new methods while minimizing the duplication of code and effort.

Due to Bragg’s law, crystallography data is inherently tabular with each observed reflection described by a Miller index. This property underlies many of the file formats for storing diffraction data; integrated intensities and any reflection-specific metadata are stored with the associated Miller index (see Fig. 1). For data analysis in Python, tabular data is commonly represented using the pandas software library [11]. `pandas.DataFrame` objects provide support for the arbitrary manipulation of tabular data, storage of heterogeneous data types, and easy integration with any scientific computing or machine learning library that supports NumPy arrays [12].

**Figure 1:**
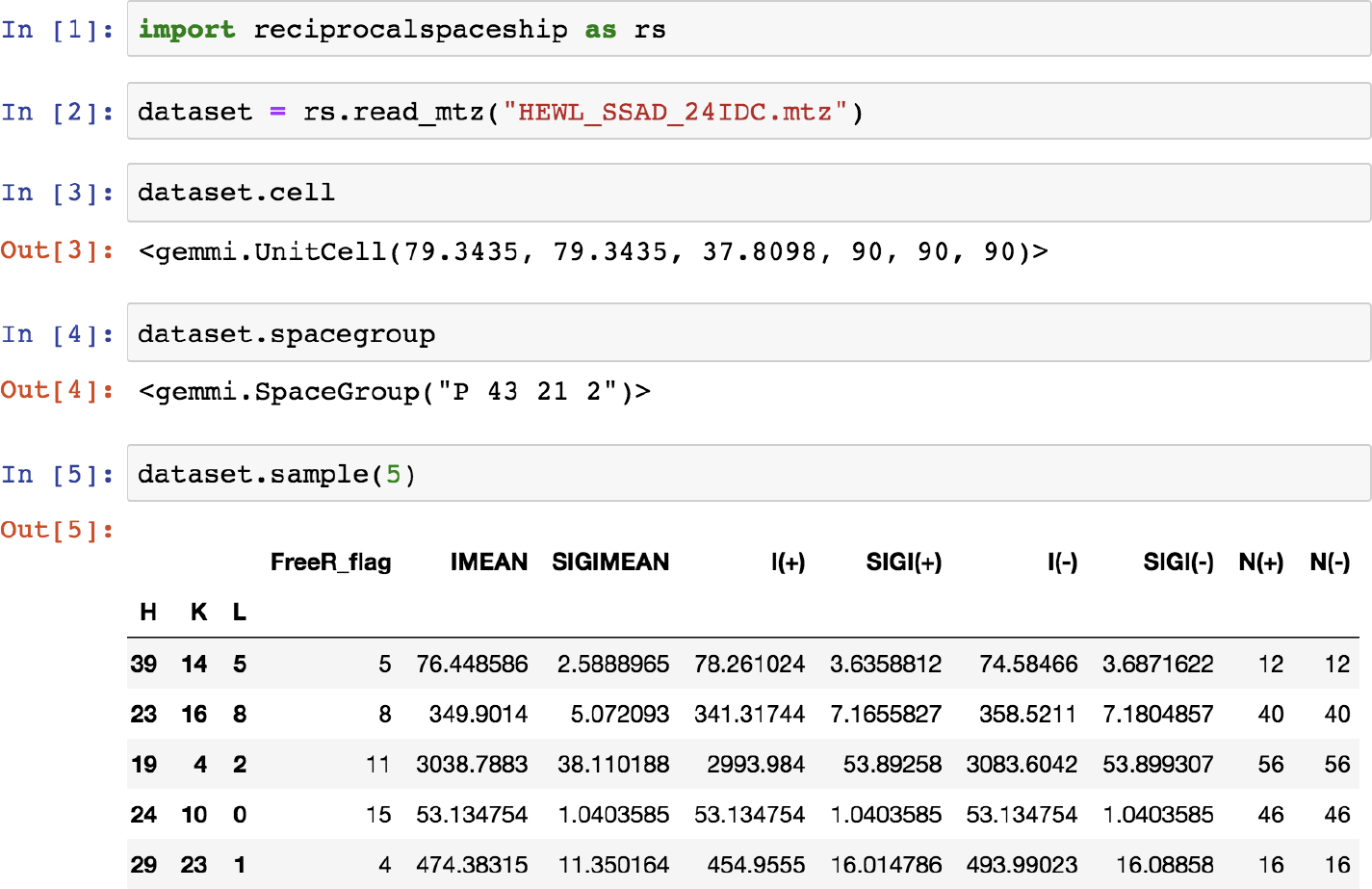
Screenshot demonstrating the use of reciprocalspaceship in a Jupyter Notebook [13]. `DataSet` objects can be used to represent reflection data with associated unit cell and space group information.

Due to the tabular nature of reflection data and the widespread use of pandas in data science, we sought to develop a library that extended the `DataFrame` for crystallographic data by providing built-in support for space groups, unit cells, and symmetry operations. This library, `reciprocalspaceship`, can be used to inspect reflection data, develop new crystallographic methods, and release reproducible analysis pipelines for X-ray diffraction experiments.

## 2 reciprocalspaceship **Library**

### 2.1 Mission Statement

`reciprocalspaceship` is a free and open-source software library with the primary goal of simplifying the analysis of crystallography data in Python. To achieve this goal, we sought to design a software library that is intuitive for both crystallographers and Python programmers. This requires full support for common crystallographic operations, as well as easy integration with the scientific computing and machine learning libraries that are developed and maintained by the Python community.

### 2.2 Design

The `DataFrame` is the core abstraction in `pandas. reciprocalspaceship` provides a `DataSet` class which extends the `DataFrame`, augmenting it to represent reflection data from X-ray diffraction experiments. `DataSet` objects store reflection data, along with the associated space group and unit cell, and can be initialized from common reflection file formats such as MTZ files (Fig. 1). By extending the `pandas DataFrame`, it is possible to preserve its core functionality while adding built-in methods to support common crystallographic operations. These operations use the GEMMI library to represent space groups and unit cells [10], and have been vectorized to increase performance.

To support compatibility with MTZ files, `reciprocalspaceship` provides custom datatypes to represent different crystallographic observables. To ensure maximum compatibility with other Python libraries, these datatypes are all represented internally using NumPy arrays of either 32-bit integers or floating-point values. Methods are also provided for inferring relevant datatypes based on the column labels used to describe the data. `DataSet` objects can contain any datatype supported by pandas, including generic Python objects.

### 2.3 Features

The primary capabilities of `reciprocalspaceship` are provided through the `DataSet` object, which builds on the core features of the pandas `DataFrame` to provide crystallographic support. These objects can represent both merged and unmerged reflection data, and provide attributes and methods that enable crystallographic data analysis. These features are summarized in Table 1.

**Table 1:**
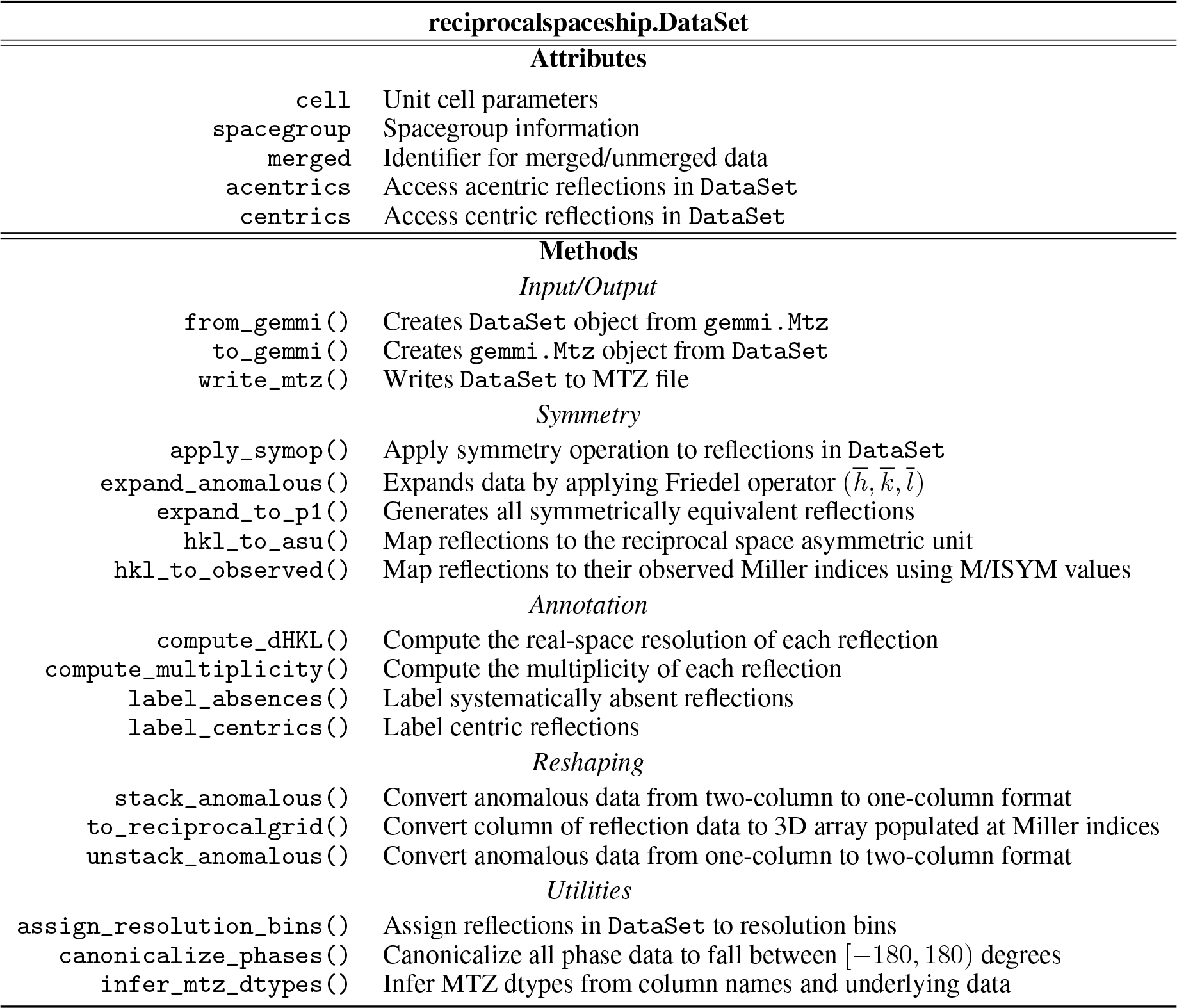
Core Features of DataSet objects

In addition to the `DataSet` object, `reciprocalspaceship` provides several algorithms that can be used for analysis. These include `merge()`, which implements the averaging of unmerged reflection data using maximum-likelihood weights, and `scale_merged_intensities()`, which implements French-Wilson scaling to account for negative merged intensities [14]. These implementations can serve as templates for the development of new analysis methods using `reciprocalspaceship`. The set of algorithms offered through this library will continue to expand as users implement new analyses intended for broader adoption.

### 2.4 Development and Documentation

`reciprocalspaceship` is maintained on GitHub to foster community involvement in its maintenance, testing, and documentation. Every change to the source code is tested using an automated suite in order to support continuous integration [15]. `reciprocalspaceship` is available through the Python Package Index (PyPI), and can be installed on most systems using `pip`. Documentation is automatically generated from the `reciprocalspaceship` GitHub repository to ensure up-to-date information is available for users. The website also includes a User Guide section describing the design and features of `reciprocalspaceship`, and examples that use the library for crystallographic applications. By committing to an open-source development model, it will be possible to maintain this library to meet the needs of crystallographers.

## 3 Examples

The following examples demonstrate the use of `reciprocalspaceship` in the analysis of crystallographic data. These examples cover the merging of scaled observed intensities, analyzing anomalous differences from a single-wavelength anomalous dispersion (SAD) experiment, and applying weights to a time-resolved difference map. These examples are intended to illustrate the breadth of crystallographic problems that can be addressed using this library, as well as its seamless integration with common scientific computing libraries. The examples are available as interactive Jupyter notebooks^1^ in the `reciprocalspaceship` documentation [13].

### 3.1 Assessing Uncertainty in Merging Statistics

Merging statistics are useful for assessing the internal consistency of a data set, and many different metrics have been proposed over the years [16, 17]. Although merging statistics are commonly reported by data reduction pipelines, they are often not reported with uncertainties and do not always give access to their underlying parameters, such as the number of resolution bins or the type of correlation coefficients to report. By facilitating inspection of the underlying reflection data, `reciprocalspaceship` can be used to write quality control scripts for automating analysis pipelines, or, as shown here, in the exploratory analysis of the properties of a single data set. By enabling crystallographers to try new statistical routines, `reciprocalspaceship` may help in the development of more robust indicators of data quality.

To illustrate this, we computed *CC*_1/2_ and *CC_anom_* for scaled, unmerged reflection data. The data were collected on a tetragonal crystal of hen egg-white lysozyme at ambient temperature and 6.5 keV. The integrated intensities were scaled in AIMLESS, and the data contains sufficient anomalous signal from the native sulfur atoms to determine experimental phases by the SAD method [18, 1, 19, 20]. Using `reciprocalspaceship`, it is possible to implement a function that merges redundant observations using inverse-variance weights in about 10 lines of code (Fig. 2a). This code takes advantage of the `groupby()` functionality inherited from `Pandas` in order to efficiently perform calculations on a per-reflection basis [11]. By randomly splitting the observed reflections by image, this function can be used to independently merge different sets of observations for computing *CC*_1/2_ and *CC_anom_*. Due to the modularity of this workflow, it is possible to repeat the random partitioning of observations to generate uncertainty estimates, and to repeat these calculations using both Pearson and Spearman correlation coefficients.

**Figure 2:**
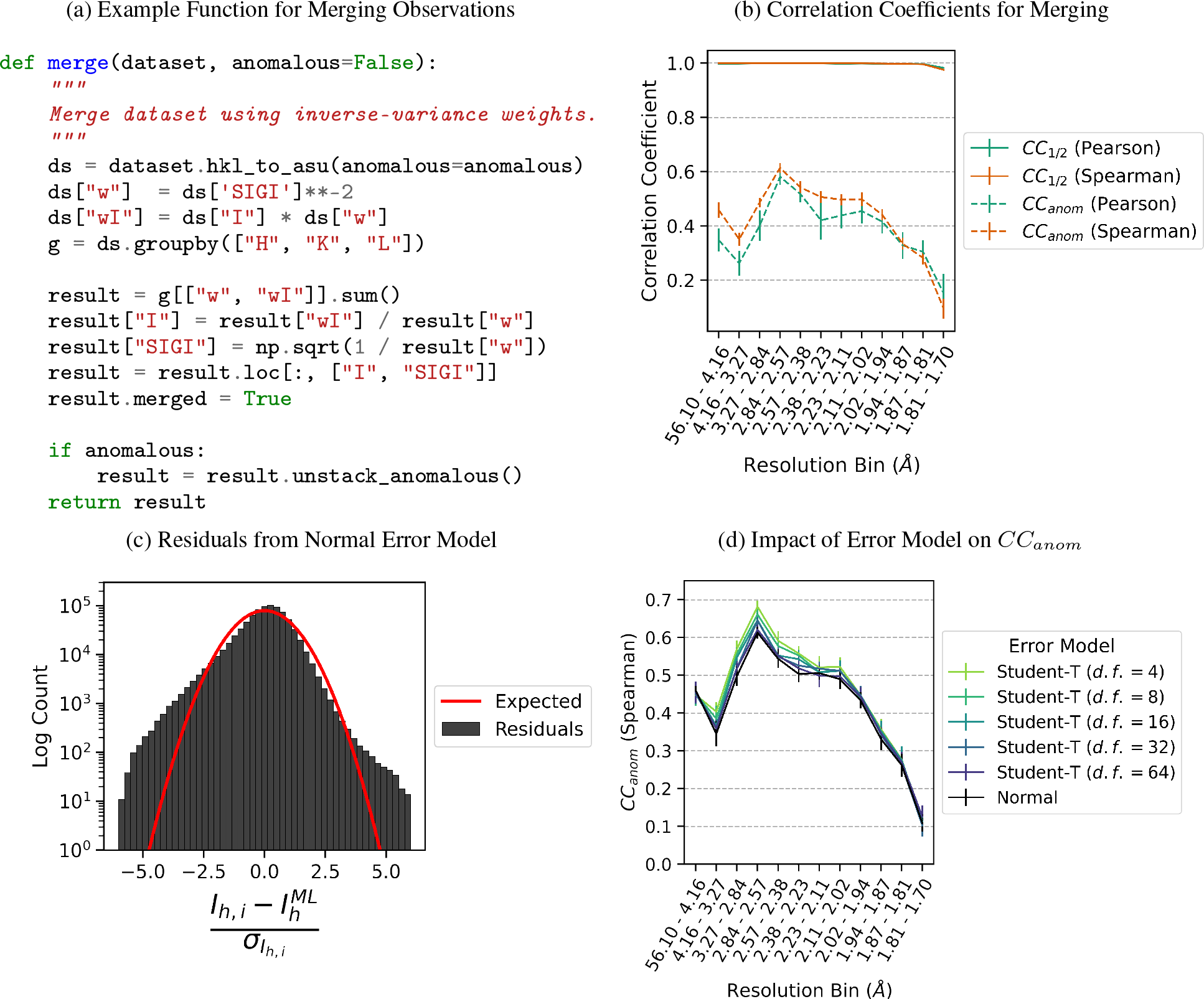
Merging statistics for a hen egg-white lysozyme sulfur SAD data set. (a) Python function for applying inverse-variance weights to obtain maximum-likelihood merged intensity estimates using `reciprocalspaceship`. (b) Correlation coefficients, *CC*_1/2_ and *CC_anom_*, from repeated two-fold cross-validation. The Pearson *CC_anom_* is more impacted by outlier measurements in low- and intermediate resolution bins than the Spearman *CC_anom_*. (c) The distribution of residuals from observed intensities differs from the expected distribution of residuals for normally distributed observations. (d) *CC_anom_* from two-fold cross-validation using Student’s t-distributed error models with varying degrees of freedom. Error models with heavier tails show improvements in *CC_anom_*. Error bars depict the mean ± standard deviation from 15 repeats of two-fold cross-validation.

As shown in Fig. 2b, high *CC*_1/2_ values indicate that the data were significantly edge-limited, which is common for data collected at low energy on strongly diffracting crystals. The *CC_anom_* values show that significant anomalous signal was obtained up to the highest resolution bin. Furthermore, the Spearman correlation coefficients are systematically higher and have smaller uncertainties in the low and intermediate resolution range suggesting the presence of outliers in the data.

### 3.2 Merging Observations with a Robust Error Model

The difference observed for *CC_anom_* between the Pearson and Spearman correlation coefficients in Fig. 2b suggests the presence of outlier observations despite the outlier rejection applied by AIMLESS [19]. Since AIMLESS assumes a normally distributed error model for its observations, such outliers can have a large impact on the estimate of the true merged intensity. We can evaluate whether a normally distributed error model is appropriate based on the distribution of residuals between the observed intensities and the estimate of the true mean. This histogram can be made in just a few lines of Python by taking advantage of the `Pandas` indexing (Fig. 2c). Compared with the expected distribution of residuals for normally distributed observations, this data set has significantly heavier tails, with many observations several standard deviations away from the merged intensity.

The residuals in Fig. 2c suggest that merging may be improved by a more robust error model that can tolerate outliers. One popular choice of robust error model is the Student’s *t*-distribution. This distribution is parameterized by a location, scale, and number of degrees of freedom, *ν*, which controls the robustness of the distribution to outliers. Importantly, the distribution approaches the normal distribution as *ν* approaches infinity. Unlike the normal distribution, there is not an analytical expression for the maximum-likelihood estimator of the true mean given a set of observations under a Student’s *t*-distributed error model. However, we can construct a simple optimization problem to recover maximum-likelihood estimates of the merged intensity for each miller index. To begin with, we write the likelihood function, which is the probability of the data as a function of the mean intensity for each miller index:

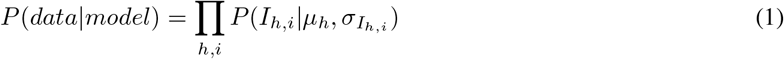

The error model *P* can be any suitable location-scale family distribution. This likelihood function asserts that the observed intensity is drawn from a distribution centered at the merged intensity, *μ_h_*, with a scale determined by the empirical standard deviation of the observation, *σ_I_h,i__*:

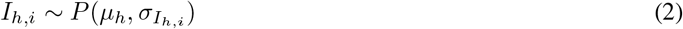

In order to recover maximum-likelihood estimates, we need only maximize equation 1 with respect to the merged intensities which are the optimization variables in this problem. Equivalently, we may minimize the negative logarithm of the likelihood:

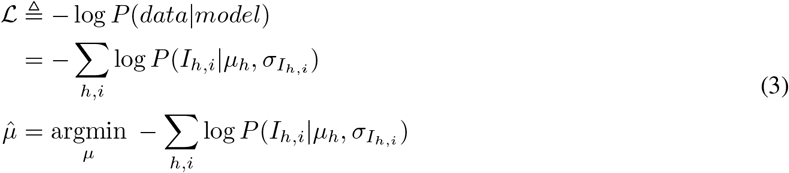

which has the advantage of converting a numerically unstable product into a sum.

This optimization was implemented in `PyTorch` in a general form that could flexibly accept a location-scale family distribution to use as an error model [21]. The data was merged using Student’s *t*-distributions with varying degrees of freedom as error models, and the resulting *CC_anom_* were compared with the normally distributed error model. The error models for fewer degrees of freedom outperformed the error models for more degrees of freedom (larger *ν*), with their performance trending towards that of the normally distributed error model (Fig. 2d).

This example demonstrates the use of `reciprocalspaceship` to construct a flexible merging function using a machine learning library. This greatly reduces the overhead required to prototype a new analysis method by making it easy to use existing and well-supported libraries. Furthermore, the benefits of using robust statistical estimators, as demonstrated by the improved *CC_anom_* values in figures 2b and 2d, suggest new avenues for improving the existing crystallographic analysis infrastructure. One such project, `careless`, is combining `reciprocalspaceship` and `TensorFlow` to use approximate Bayesian inference in order to develop new scaling and merging routines [22, 23].

### 3.3 Revisiting French-Wilson Scaling

In the previous example, we identified anomalous differences from a room-temperature sulfur SAD experiment. Here, we will examine this anomalous signal in real space by making an anomalous difference map. Before we can make a map, it is necessary to scale the merged intensities to account for any negative values that may result from background subtraction during integration. This is commonly handled using a Bayesian approach first proposed by French and Wilson [14]. Briefly, this algorithm works by solving an integral:

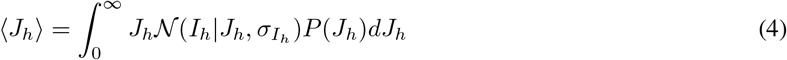

where the likelihood, 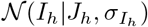, is taken to be normally distributed with the empirical standard deviation. The prior distribution, *P*(*J_h_*), is the Wilson distribution:

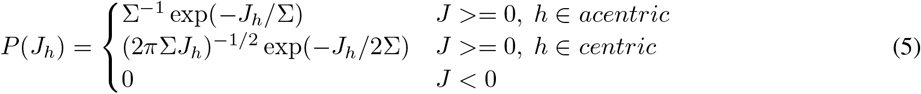

which is parameterized by Σ, the mean intensity of reflections at the appropriate resolution. In order to estimate Σ for each reflection, the classic French-Wilson scaling algorithm computes the mean intensity of reflections in resolution shells, and interpolates the mean values from shells adjacent to the particular reflection. Since the functional form of the prior distribution has strictly positive support [24], the expectations computed from equation 4 are necessarily positive. Furthermore, the posterior structure factor amplitudes can be estimated as part of the same subroutine using the following integral:

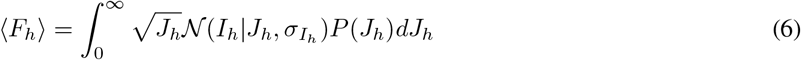

This scaling method is implemented in `reciprocalspaceship` as `scale_merged_intensities()`, though this implementation differs significantly from the classical one in several regards. Notably, rather than computing mean values in shells, we use a Gaussian smoother [25, chapter 14.7.4-5] to regress the mean of the intensity distributions, Σ, against resolution. This regression model is quite flexible and offers an anisotropic mode which estimates the mean intensity locally as a function of the Miller indices. Whereas the original paper computed the posterior by interpolating a table of cached results [14], our implementation uses Chebyshev-Gauss quadrature to evaluate the integrals on the fly. We generate quadrature points and weights with `NumPy` [12] and compute the relevant log probabilities using the distribution classes implemented in SciPy [26]. Our implementation is tested for consistency with the original paper [14] and with `CCTBX`[4].

The merged intensities from the sulfur SAD experiment were rescaled and converted to structure factors using `scale_merged_intensities()`. This operation leaves large intensities relatively unchanged, while ensuring that any negative values become strictly positive (Fig. 3a). Anomalous differences of the structure factor amplitudes were computed between Friedel pairs. The anomalous difference map shown in Fig. 3b was then constructed using phases derived from the refined model (PDB: 7L84). The map shows significant anomalous peaks at a 5*σ* contour, with the density localized to each of the 10 sulfur atoms in the lysozyme structure.

**Figure 3:**
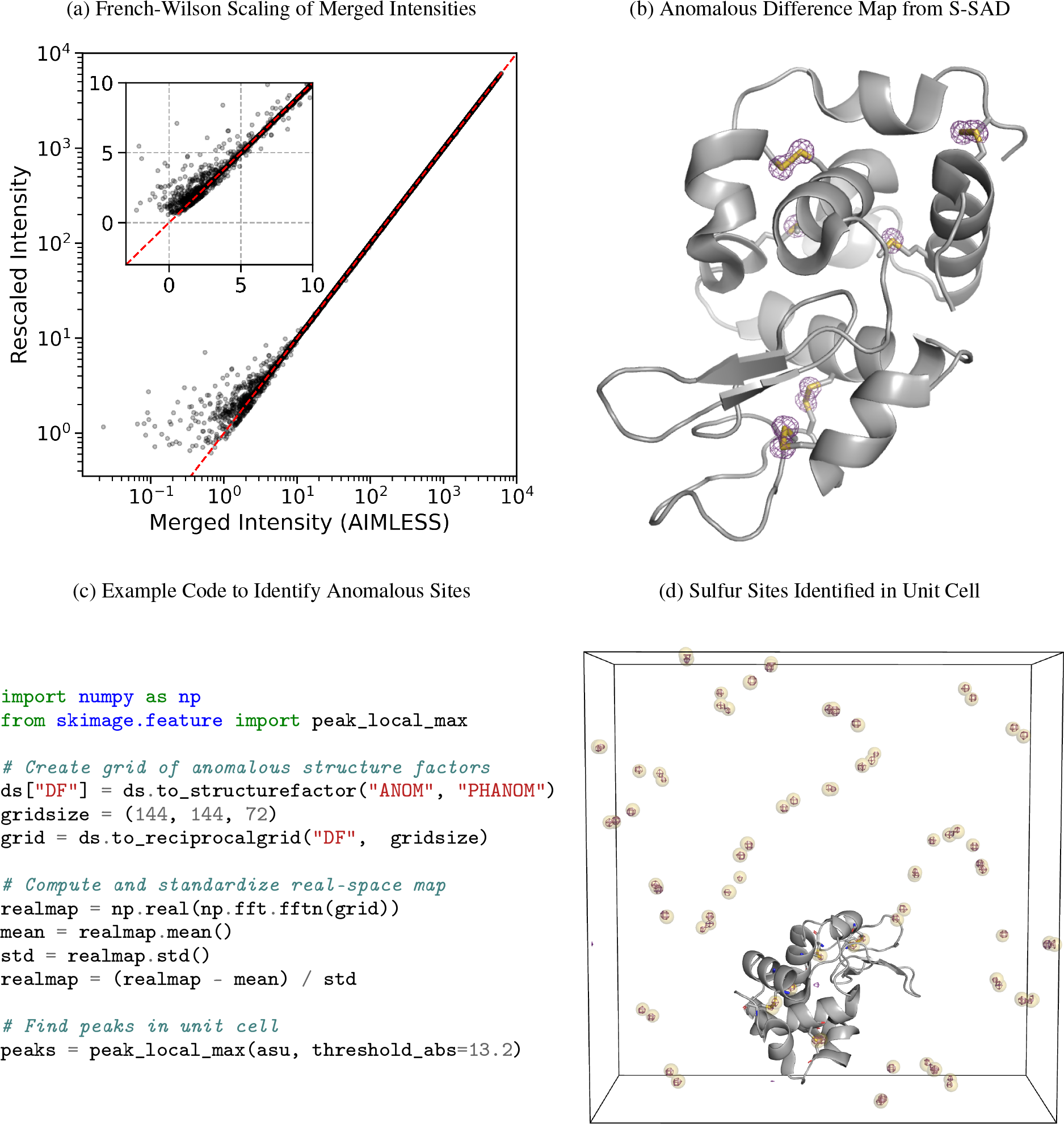
Analysis of anomalous differences from a sulfur SAD experiment. (a) French-Wilson scaling of merged intensities. Large intensities are relatively unchanged, while small and negative intensities are rescaled to be strictly positive. The red dashed line shows *y = x*, and the inset highlights the small and negative merged intensities. (b) Anomalous difference map using difference structure factor amplitudes derived from room-temperature sulfur SAD data set and phases from the refined model (PDB: 7L84). Map is contoured at 5*σ*. (c) Example code for identifying anomalous scattering sites in an anomalous difference map. (d) Unit cell containing sulfur sites identified using the code snippet (yellow spheres), overlaid with the anomalous difference map (contoured at 10*σ*). The protein molecule from one asymmetric unit of PDB: 7L84 is shown in gray.

### 3.4 Identifying Anomalous Scattering Atoms in Real Space

The anomalous difference map shown in Fig. 3b was rendered in PyMOL (Schrödinger, LLC) from the anomalous difference amplitudes and phases. It is also possible to compute a real-space map using `reciprocalspaceship` and NumPy, which enables one to use image processing software to automate the identification of anomalous scattering atoms. This process is illustrated in Fig. 3c. This code snippet arranges the complex anomalous structure factors on a reciprocal space grid, and then computes the real-space anomalous difference map using the Fast Fourier transform [27] function in NumPy [12]. `scikit-image`, an image processing library [28], can be used to identify peaks in the map. This procedure successfully identifies the 80 sulfur sites in the tetragonal lysozyme unit cell (10 sulfurs per copy, 8 copies). The automatically identified sites are overlaid with the anomalous difference map in Fig. 3d.

This example illustrates the use of `reciprocalspaceship` to produce real-space maps from structure factors. Importantly, due to the seamless integration with NumPy, it is possible to take advantage of Python image processing libraries for identifying peaks in the real-space density. Due to the wealth of libraries and tools written by the Python community, this feature of `reciprocalspaceship` can provide the opportunity to develop and test new algorithms rapidly. In this manner, the use of `reciprocalspaceship` could simplify existing data processing pipelines, and perhaps be useful in the development of new methods in crystallographic data analysis or structural bioinformatics.

### 3.5 Applying Weights to a Time-Resolved Difference Map

Time-resolved crystallography experiments make use of X-ray diffraction to monitor structural changes in a crystalline sample. Commonly, structural changes are initially evaluated on the basis of isomorphous difference maps. Such maps are computed by estimating the difference in structure factor amplitudes of the sample before and after a perturbation, such as a laser pulse. Combining these |*F_on_*| – |*F_off_*| differences with ground state phases from a reference structure yields an estimate of the differences between the electron density of the sample before and after the perturbation. Difference maps are often noisy due to systematic errors or scaling artifacts, and are frequently weighted by the magnitude of the difference signal and/or the error estimates associated with the empirical differences in structure factor amplitudes. In this example we will visualize the effects of applying weights to a time-resolved difference map of photoactive yellow protein (PYP). PYP is a model system in time-resolved crystallography due to the trans-to-cis isomerization of its 4-hydroxycinnamyl chromophore which occurs upon absorption of blue light [29]. This data set was collected at the BioCARS Laue beamline APS-14-ID, and is composed of matched images collected in the dark and 2ms after illumination with blue light. This data was collected and provided by Marius Schmidt and Vukica Šrajer. Several schemes have been used to apply weights to time-resolved difference maps. Many of them take the form of Equation 7, involving a term based on the uncertainty in the difference structure factor amplitude (*σ*_Δ*F*_) and optionally, a scale term based on the the magnitude of the observed difference structure factor amplitude (|Δ*F*|):

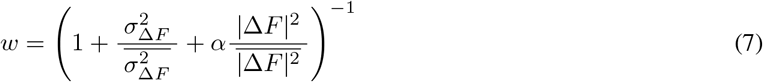

With *α* = 0, these weights take the form derived by Ursby and Bourgeois [30]. The Δ*F*-dependent term downweights the influence of outliers in the data set resulting from poorly measured differences by assigning lower weights to their map coefficients. The degree of skepticism about large differences is controlled by the *α* parameter. *α* values of 1.0 [31] and 0.05 [8] have been reported in the literature.

The weighting function given by Equation 7 can be expressed in a few lines of Python that apply weights based on the values of |Δ*F*| and *σ*_Δ*F*_ in an `rs.DataSet` object (Fig. 4a). The weights computed for the PYP data set are illustrated in Fig. 4b. Difference structure factors with low signal-to-noise ratios (large *σ*_Δ*F*_ relative to |Δ*F*|) or large difference structure factor amplitudes are assigned lower weight. The unweighted and weighted difference maps were then made using phases derived from the ground-state model (PDB: 2PHY). The side-by-side comparison of these difference maps shows that the weights greatly improve the interpretation of the structural changes—emphasizing the trans-to-cis isomerization of the chromophore as well as concerted changes in the nearby Arg52 and Phe96 sidechains (Fig. 4c and 4d).

**Figure 4:**
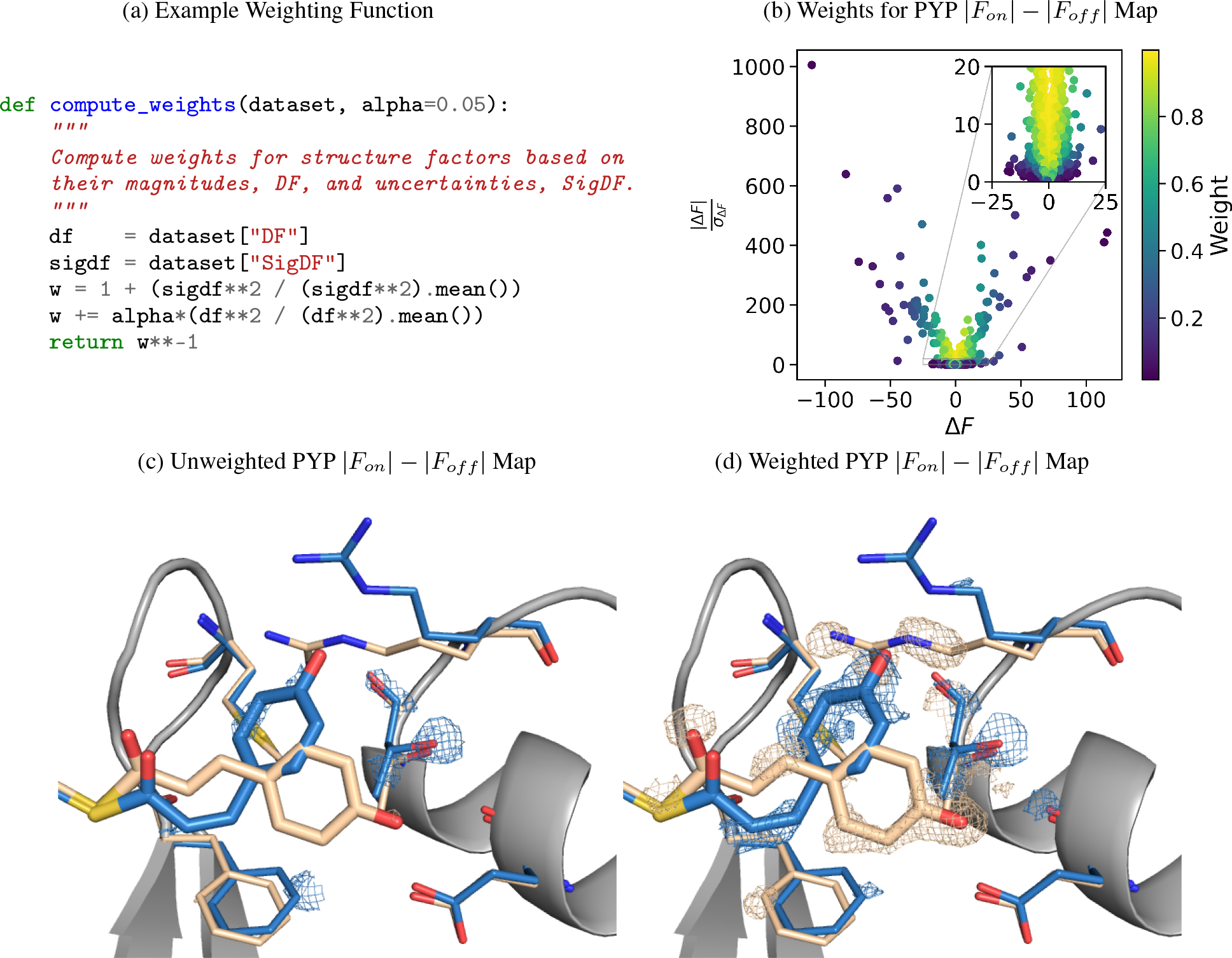
Weighting a time-resolved difference map. (a) Python function for applying weights to arrays of difference structure factor amplitudes and uncertainties. (b) Scatter plot showing the weights assigned to each observed difference structure factor amplitude with *α* = 0.05. (c) Unweighted PYP |*F_on_* | — |*F_off_*| difference map in the vicinity of the PYP chromophore. (d) Weighted PYP |*F_on_*| — |*F_off_*| difference map with *α* = 0.05. The trans (ground state) PYP structure (tan) is taken from PDB: 2PHY, and the cis (excited, pB state) PYP structure (blue) is taken from PDB: 3UME. The difference maps are contoured at ±3σ.

This example illustrates the use of `reciprocalspaceship` for creating custom maps. Importantly, it demonstrates both the exploratory analysis of different weighting schemes, as well as writing MTZ files including different weight columns. These can be used to visualize the impact of the different weights in a molecular visualization suite.

## 4 Discussion

`reciprocalspaceship` is a Python library that can form the foundation for the development of new methods in crystallographic data analysis. This library provides a `DataSet` object that can conveniently represent tabular reflection data while adhering to common practices in Python data analysis. This empowers crystallographers to write idiomatic Python code to analyze their experiments while having full support for the necessary features of crystallographic analysis, such as symmetry operations, unit cells, and spacegroups. Example applications were presented which use this library for merging scaled reflections, analyzing anomalous differences from a SAD experiment, and for observing the impact of weights on a time-resolved difference map. These examples illustrate how `reciprocalspaceship` could be used in several different contexts, producing useful analyses with relatively short scripts and functions that can take full advantage of the existing Python ecosystem.

`reciprocalspaceship` can be used for exploratory data analysis – allowing one to inspect interesting properties of an important data set. Or it can be used to prototype, develop and ship new methods and algorithms for analyzing data sets [22]. Furthermore, this library can be useful in teaching crystallography by allowing students to familiarize themselves with reflection data, space groups and symmetry, and the implementation of commonly-used algorithms. This library lowers the barrier to entry for crystallographic software development by using a framework familiar to Python data scientists.

## 5 Data and Code Availability

`reciprocalspaceship` and worked-out examples are available on GitHub at https://github.com/Hekstra-Lab/reciprocalspaceship, and can be installed directly from the Python Package Index (PyPI). The code used in these examples are available in the `reciprocalspaceship` documentation, and the interactive Jupyter notebooks and all supporting data can be downloaded directly from the Examples directory of the GitHub repository.

## 6 Acknowledgements

We thank the staff at the Northeastern Collaborative Access Team (NE-CAT), beamline 24-ID-C of the Advanced Photon Source, for supporting our room-temperature crystallography experiments, with special thanks to Igor Kourinov. NE-CAT beamlines are supported by the National Institute of General Medical Sciences, NIH (P30 GM124165), using resources of the Advanced Photon Source, a U.S. Department of Energy (DOE) Office of Science User Facility operated for the DOE Office of Science by Argonne National Laboratory under Contract No. DE-AC02-06CH11357. We also thank Marius Schmidt and Vukica Šrajer for the time-resolved Laue diffraction data of photoactive yellow protein. This work was supported by the Searle Scholarship Program (SSP-2018-3240) and a fellowship from the George W. Merck Fund of the New York Community Trust (338034). J.B.G. was supported by the National Science Foundation Graduate Research Fellowship under Grant No. DGE1745303.

## Footnotes

1 https://mybinder.org/v2/gh/Hekstra-Lab/reciprocalspaceship/master?filepath=docs%2Fexamples

## References

[1] Paul D. Adams, Pavel V. Afonine, Gábor Bunkóczi, Vincent B. Chen, Ian W. Davis, Nathaniel Echols, Jeffrey J. Headd, Li-Wei Hung, Gary J. Kapral, Ralf W. Grosse-Kunstleve, Airlie J. McCoy, Nigel W. Moriarty, Robert Oeffner, Randy J. Read, David C. Richardson, Jane S. Richardson, Thomas C. Terwilliger, and Peter H. Zwart. *PHENIX*: a comprehensive Python-based system for macromolecular structure solution. Acta Crystallographica Section D, 66(2):213–221, Feb 2010.

[2] Martyn D. Winn, Charles C. Ballard, Kevin D. Cowtan, Eleanor J. Dodson, Paul Emsley, Phil R. Evans, Ronan M. Keegan, Eugene B. Krissinel, Andrew G. W. Leslie, Airlie McCoy, Stuart J. McNicholas, Garib N. Murshudov, Navraj S. Pannu, Elizabeth A. Potterton, Harold R. Powell, Randy J. Read, Alexei Vagin, and Keith S. Wilson. Overview of the CCP4 suite and current developments. Acta Crystallographica Section D, 67(4):235–242, Apr 2011.

[3] Graeme Winter, David G. Waterman, James M. Parkhurst, Aaron S. Brewster, Richard J. Gildea, Markus Gerstel, Luis Fuentes-Montero, Melanie Vollmar, Tara Michels-Clark, Iris D. Young, Nicholas K. Sauter, and Gwyndaf Evans. *DIALS*: implementation and evaluation of anew integration package. Acta Crystallographica Section D, 74(2):85–97, Feb 2018.

[4] Ralf W. Grosse-Kunstleve, Nicholas K. Sauter, Nigel W. Moriarty, and Paul D. Adams. The *Computational Crystallography Toolbox*: crystallographic algorithms in a reusable software framework. Journal of Applied Crystallography, 35(1):126–136, Feb 2002.

[5] Wolfgang Kabsch. XDS. Acta Crystallographica Section D, 66(2):125–132, Feb 2010.

[6] Wolfgang Kabsch. Integration, scaling, space-group assignment and post-refinement. Acta Crystallographica Section D, 66(2):133–144, Feb 2010.

[7] Zbyszek Otwinowski and Wladek Minor. Processing of x-ray diffraction data collected in oscillation mode. In Macromolecular Crystallography Part A, volume 276 of Methods in Enzymology, pages 307 – 326. Academic Press, 1997.

[8] Doeke R. Hekstra, K. Ian White, Michael A. Socolich, Robert W. Henning, Vukica Šrajer, and Rama Ranganathan. Electric-field-stimulated protein mechanics. Nature, 540(7633):400–405, 2016.

[9] Robert Dods, Petra Båth, Dmitry Morozov, Viktor Ahlberg Gagnér, David Arnlund, Hoi Ling Luk, Joachim Kübel, Michał Maj, Adams Vallejos, Cecilia Wickstrand, Robert Bosman, Kenneth R Beyerlein, Garrett Nelson, Mengning Liang, Despina Milathianaki, Joseph Robinson, Rajiv Harimoorthy, Peter Berntsen, Erik Malmerberg, Linda Johansson, Rebecka Andersson, Sergio Carbajo, Elin Claesson, Chelsie E Conrad, Peter Dahl, Greger Hammarin, Mark S Hunter, Chufeng Li, Stella Lisova, Antoine Royant, Cecilia Safari, Amit Sharma, Garth J Williams, Oleksandr Yefanov, Sebastian Westenhoff, Jan Davidsson, Daniel P DePonte, Sébastien Boutet, Anton Barty, Gergely Katona, Gerrit Groenhof, Gisela Brändén, and Richard Neutze. Ultrafast structural changes within a photosynthetic reaction centre. Nature, 589(7841):310–314, 2021.

[10] CCP4 and Global Phasing Ltd. Gemmi – library for structural biology [software]. https://github.com/project-gemmi/gemmi, 2020.

[11] Jeff Reback, Wes McKinney, jbrockmendel, Joris Van den Bossche, Tom Augspurger, Phillip Cloud, gfyoung, Simon Hawkins, Sinhrks, Matthew Roeschke, Adam Klein, Terji Petersen, Jeff Tratner, Chang She, William Ayd, Shahar Naveh, Marc Garcia, Jeremy Schendel, Andy Hayden, Daniel Saxton, patrick, Vytautas Jancauskas, Ali McMaster, Pietro Battiston, Skipper Seabold, Marco Gorelli, Kaiqi Dong, chris b, h vetinari, and Stephan Hoyer. pandas-dev/pandas: Pandas 1.2.1 [software]. https://doi.org/10.5281/zenodo.3509134, January 2021.

[12] Charles R Harris, K Jarrod Millman, Stéfan J van der Walt, Ralf Gommers, Pauli Virtanen, David Cournapeau, Eric Wieser, Julian Taylor, Sebastian Berg, Nathaniel J Smith, Robert Kern, Matti Picus, Stephan Hoyer, Marten H van Kerkwijk, Matthew Brett, Allan Haldane, Jaime Fernández del Río, Mark Wiebe, Pearu Peterson, Pierre Gérard-Marchant, Kevin Sheppard, Tyler Reddy, Warren Weckesser, Hameer Abbasi, Christoph Gohlke, and Travis E Oliphant. Array programming with NumPy. Nature, 585(7825):357–362, 2020.

[13] Thomas Kluyver, Benjamin Ragan-Kelley, Fernando Pérez, Brian Granger, Matthias Bussonnier, Jonathan Frederic, Kyle Kelley, Jessica Hamrick, Jason Grout, Sylvain Corlay, Paul Ivanov, Damián Avila, Safia Abdalla, Carol Willing, and Jupyter development team. Jupyter notebooks – a publishing format for reproducible computational workflows. In Fernando Loizides and Birgit Scmidt, editors, Positioning and Power in Academic Publishing: Players, Agents and Agendas, pages 87–90, Netherlands, 2016. IOS Press.

[14] S. French and K. Wilson. On the treatment of negative intensity observations. Acta Crystallographica Section A, 34(4):517–525, Jul 1978.

[15] Holger Krekel, Bruno Oliveira, Ronny Pfannschmidt, Floris Bruynooghe, Brianna Laugher, and Florian Bruhin. pytest 6.2.1 [software]. https://github.com/pytest-dev/pytest, 2020.

[16] Manfred S. Weiss. Global indicators of X-ray data quality. Journal of Applied Crystallography, 34(2):130–135, Apr 2001.

[17] P Andrew Karplus and Kay Diederichs. Linking crystallographic model and data quality. Science, 336(6084):1030–1033, 2012.

[18] Jack B. Greisman, Kevin M. Dalton, and Doeke R. Hekstra. Hen Egg White Lysozyme by Native S-SAD at Room Temperature (Version 1.0.0) [Dataset]. https://doi.org/10.5281/zenodo.4426679, January 2021.

[19] Philip R. Evans and Garib N. Murshudov. How good are my data and what is the resolution? Acta Crystallographica Section D, 69(7):1204–1214, Jul 2013.

[20] T. C. Terwilliger, P. D. Adams, R. J. Read, A. J. McCoy, N. W. Moriarty, R. W. Grosse-Kunstleve, P V. Afonine, P. H. Zwart, and L.-W. Hung. Decision-making in structure solution using Bayesian estimates of map quality: the PHENIX AutoSol wizard. Acta Crystallographica Section D: Biological Crystallography, 65(6):582–601, June 2009.

[21] Adam Paszke, Sam Gross, Francisco Massa, Adam Lerer, James Bradbury, Gregory Chanan, Trevor Killeen, Zeming Lin, Natalia Gimelshein, Luca Antiga, Alban Desmaison, Andreas Kopf, Edward Yang, Zachary DeVito, Martin Raison, Alykhan Tejani, Sasank Chilamkurthy, Benoit Steiner, Lu Fang, Junjie Bai, and Soumith Chintala. Pytorch: An imperative style, high-performance deep learning library. In H. Wallach, H. Larochelle, A. Beygelzimer, F. d’Alché-Buc, E. Fox, and R. Garnett, editors, Advances in Neural Information Processing Systems 32, pages 8024–8035. Curran Associates, Inc., 2019.

[22] Kevin M. Dalton, Jack B. Greisman, and Doeke R. Hekstra. Careless: A variational bayesian model for merging x-ray diffraction data. bioRxiv, 2021.

[23] Martín Abadi, Ashish Agarwal, Paul Barham, Eugene Brevdo, Zhifeng Chen, Craig Citro, Greg S. Corrado, Andy Davis, Jeffrey Dean, Matthieu Devin, Sanjay Ghemawat, Ian Goodfellow, Andrew Harp, Geoffrey Irving, Michael Isard, Yangqing Jia, Rafal Jozefowicz, Lukasz Kaiser, Manjunath Kudlur, Josh Levenberg, Dandelion Mané, Rajat Monga, Sherry Moore, Derek Murray, Chris Olah, Mike Schuster, Jonathon Shlens, Benoit Steiner, Ilya Sutskever, Kunal Talwar, Paul Tucker, Vincent Vanhoucke, Vijay Vasudevan, Fernanda Viégas, Oriol Vinyals, Pete Warden, Martin Wattenberg, Martin Wicke, Yuan Yu, and Xiaoqiang Zheng. TensorFlow: Large-scale machine learning on heterogeneous systems, 2015. Software available from tensorflow.org.

[24] A. J. C. Wilson. The probability distribution of X-ray intensities. Acta Crystallographica, 2(5):318–321, October 1949. Number: 5 Publisher: International Union of Crystallography.

[25] Kevin P. Murphy. Machine Learning: A Probabilistic Perspective. MIT Press, Cambridge, United States, 2012.

[26] Pauli Virtanen, Ralf Gommers, Travis E Oliphant, Matt Haberland, Tyler Reddy, David Cournapeau, Evgeni Burovski, Pearu Peterson, Warren Weckesser, Jonathan Bright, Stéfan J van der Walt, Matthew Brett, Joshua Wilson, K Jarrod Millman, Nikolay Mayorov, Andrew R J Nelson, Eric Jones, Robert Kern, Eric Larson, C J Carey, ílhan Polat, Yu Feng, Eric W Moore, Jake VanderPlas, Denis Laxalde, Josef Perktold, Robert Cimrman, Ian Henriksen, E A Quintero, Charles R Harris, Anne M Archibald, Antônio H Ribeiro, Fabian Pedregosa, Paul van Mulbregt, Aditya Vijaykumar, Alessandro Pietro Bardelli, Alex Rothberg, Andreas Hilboll, Andreas Kloeckner, Anthony Scopatz, Antony Lee, Ariel Rokem, C Nathan Woods, Chad Fulton, Charles Masson, Christian Häggström, Clark Fitzgerald, David A Nicholson, David R Hagen, Dmitrii V Pasechnik, Emanuele Olivetti, Eric Martin, Eric Wieser, Fabrice Silva, Felix Lenders, Florian Wilhelm, G Young, Gavin A Price, Gert-Ludwig Ingold, Gregory E Allen, Gregory R Lee, Hervé Audren, Irvin Probst, Jörg P Dietrich, Jacob Silterra, James T Webber, Janko Slavic, Joel Nothman, Johannes Buchner, Johannes Kulick, Johannes L Schönberger, José Vinícius de Miranda Cardoso, Joscha Reimer, Joseph Harrington, Juan Luis Cano Rodríguez, Juan Nunez-Iglesias, Justin Kuczynski, Kevin Tritz, Martin Thoma, Matthew Newville, Matthias Kümmerer, Maximilian Bolingbroke, Michael Tartre, Mikhail Pak, Nathaniel J Smith, Nikolai Nowaczyk, Nikolay Shebanov, Oleksandr Pavlyk, Per A Brodtkorb, Perry Lee, Robert T McGibbon, Roman Feldbauer, Sam Lewis, Sam Tygier, Scott Sievert, Sebastiano Vigna, Stefan Peterson, Surhud More, Tadeusz Pudlik, Takuya Oshima, Thomas J Pingel, Thomas P Robitaille, Thomas Spura, Thouis R Jones, Tim Cera, Tim Leslie, Tiziano Zito, Tom Krauss, Utkarsh Upadhyay, Yaroslav O Halchenko, Yoshiki Vázquez-Baeza, and SciPy 1.0 Contributors. SciPy 1.0: fundamental algorithms for scientific computing in Python. Nature Methods, 17(3):261–272, 2020.

[27] James W. Cooley and John W. Tukey. An algorithm for the machine calculation of complex Fourier series. Mathematics of Computation, 19(90):297–301, 1965.

[28] Stéfan van der Walt, Johannes L. Schönberger, Juan Nunez-Iglesias, François Boulogne, Joshua D. Warner, Neil Yager, Emmanuelle Gouillart, Tony Yu, and the scikit-image contributors. scikit-image: image processing in Python. PeerJ, 2:e453, 6 2014.

[29] Ulrich K. Genick, Gloria E. O. Borgstahl, Kingman Ng, Zhong Ren, Claude Pradervand, Patrick M. Burke, Vukica Šrajer, Tsu-Yi Teng, Wilfried Schildkamp, Duncan E. McRee, Keith Moffat, and Elizabeth D. Getzoff. Structure of a protein photocycle intermediate by millisecond time-resolved crystallography. Science, 275(5305):1471–1475, 1997.

[30] T. Ursby and D. Bourgeois. Improved Estimation of Structure-Factor Difference Amplitudes from Poorly Accurate Data. Acta Crystallographica Section A, 53(5):564–575, Sep 1997.

[31] Vukica Šrajer, Zhong Ren, Tsu-Yi Teng, Marius Schmidt, Thomas Ursby, Dominique Bourgeois, Claude Pradervand, Wilfried Schildkamp, Michael Wulff, and Keith Moffat. Protein conformational relaxation and ligand migration in myoglobin: A nanosecond to millisecond molecular movie from time-resolved laue x-ray diffraction. Biochemistry, 40(46):13802–13815, 2001.

